# Evaluation of Sb-1 bacteriophage activity in enhancing antibiotic efficacy against biofilm, degrading the exopolysaccharide matrix and targeting persister cells of *Staphylococcus aureus*

**DOI:** 10.1101/312736

**Authors:** Tamta Tkhilaishvili, Lisa Lombardi, Ann-Brit Klatt, Andrej Trampuz, Mariagrazia Di Luca

## Abstract

Most research on phage therapy focused on planktonic bacteria, whereas bacteriophage activity against biofilms is limited. We evaluated the capability of *Staphylococcus aureus*-specific bacteriophage Sb-1 to eradicate biofilm alone and in combination with different classes of antibiotics, to degrade the extracellular matrix and target persister cells. Biofilm of methicillin-resistant *S. aureus* (MRSA) ATCC 43300 was treated with Sb-1 alone or in (simultaneous or staggered) combination with either fosfomycin, rifampin, vancomycin, daptomycin or ciprofloxacin. The matrix was visualized by confocal fluorescent microscopy. Persister cells were treated with 10^4^ and 10^7^ PFU/mL Sb-1 for 3 hours in PBS, followed by CFU counting. Alternatively, bacteria were washed and incubated in fresh BHI medium and the bacterial growth assessed after further 24-hours. Pre-treatment with Sb-1 followed by the administration of sub-inhibitory concentrations of antibiotic exerted a considerable synergistic effect in eradicating MRSA biofilm. Sb-1 determined a dose-dependent reduction of matrix exopolysaccharide. 10^7^ PFU/mL Sb-1 showed direct killing activity on persisters. However, even a lower titer had lytic activity when phage-treated persister cells were inoculated in fresh medium, reverting to a normal-growing phenotype. This study provides valuable data regarding the capability of Sb-1 to enhance antibiotic efficacy, exhibiting specific antibiofilm features. Its ability to degrade the MRSA polysaccharide matrix and target persister cells makes Sb-1 suitable for the therapy of biofilm-associated infections.

## INTRODUCTION

*Staphylococcus aureus* is a major cause of both community-and hospital-acquired infections and represents a significant burden on the healthcare system. In addition, *S. aureus* is also involved in infections of medical implants and host tissue due to its ability to form biofilms, which play an important role in the persistence of chronic infections (1, 2). A biofilm is defined as a sessile microbial community in which microorganisms live attached to a surface in a highly hydrated extracellular matrix (3), which in case of *S. aureus* is composed of host factors, secreted and lysis-derived proteins, polysaccharides, and eDNA (4). Biofilm cell population shows structural and functional heterogeneity. Depletion of nutrients causes microbes to enter a metabolically quiescent state. In addition, biofilms accommodate a high level of persister cells, an isogenic sub-population of bacteria tolerant to antibiotics that is characterized by a slow-or non-growing state (5, 6). The presence of the extracellular matrix and the heterogeneity of the cellular metabolic status make biofilm-embedded bacteria up to 1000 times more resistant to most antimicrobial agents than their planktonic counterparts (7, 8). The challenge of biofilm treatment spur scientists to investigate alternative strategies for its eradication. In this context, bacteriophage therapy, based on the administration of viruses selectively infecting target bacteria (9, 10), has re-emerged as potential therapeutic option in cases where antibiotics alone are not able to eradicate the biofilm (11, 12). Recently, different studies investigated the effect of the combined phage/antibiotic treatment against biofilm-embedded cells. Ryan et al showed that a synergistic effect of cefotaxime was observed when used in combination with the T4 phage against *Escherichia coli* biofilm (13). Two independent studies also demonstrated that pre-treating *Pseudomonas aeruginosa* (14) and *S. aureus* (15) biofilms with bacteriophages before antibiotic administration determined a major reduction of bacterial viability compared to the effect observed with the simultaneous administration of these therapeutic agents. However, none of the above-mentioned papers investigated the mechanisms through which phages can enhance antibiotics activity against biofilms.

In this study, we aimed to evaluate the susceptibility of *S. aureus* biofilm to Sb-1 bacteriophage administered with fosfomycin, rifampin, vancomycin daptomycin and ciprofloxacin, either simultaneously or in staggered treatment. In addition, we examined the ability of Sb-1 to degrade extracellular matrix and to target persister and metabolically less active cells of *S. aureus* biofilm. Sb-1 exhibits strong synergistic effect when phage administration precedes the antibiotic treatment. Moreover, Sb-1 shows specific anti-biofilm features, as - unlike most conventional antibiotics - it is able to degrade the exopolysaccharide components of the matrix and to kill persister cells. These results argue that including phages pre-treatment in the current antibiotic therapy might be an advantage for biofilm eradication.

## RESULTS

### Sb-1 phage lyses *Staphylococcus aureus* planktonic cells, but it does not eradicate the biofilm

We evaluated the lytic spectrum of Sb-1 on a collection of *S. aureus* clinical isolates from implant-associated infections (29 MSSA and 28 MRSA strains). The MRSA ATCC 43300 was also included. 27 MRSA strains – including ATCC 43300 – and 20 MSSA strains were susceptible to Sb-1, as indicated by the appearance of plaques of lysis. The parameters of the viral cycle of the phage during the infection of ATCC 43300, which was used for anti-biofilm experiments, are reported in Table 1. In order to evaluate Sb-1 potential anti-biofilm activity, 24 hour-biofilm-coated glass beads were exposed to Sb-1 titers ranging from 10^2^ to 10^7^ PFU/mL for 24 hours, and then transferred into fresh media, where the heat flow produced by the replication of cells still adhering on the bead was followed by microcalorimetry. The treatment with Sb-1 determined a strong reduction in the heat flow as compared to the growth control already at the low titers tested (Figure 1). However, only the highest titer (10^7^ PFU/mL) was able to inhibit the heat production more than 90% compared to the growth control, defined as the MBBC according to (16). Moreover, when sonication fluids obtained from the beads were plated, bacterial growth was observed, proving that, despite the strong reduction in viable cells, 10^7^ PFU/mL of Sb-1 could not completely eradicate the biofilm.

**Figure 1.**
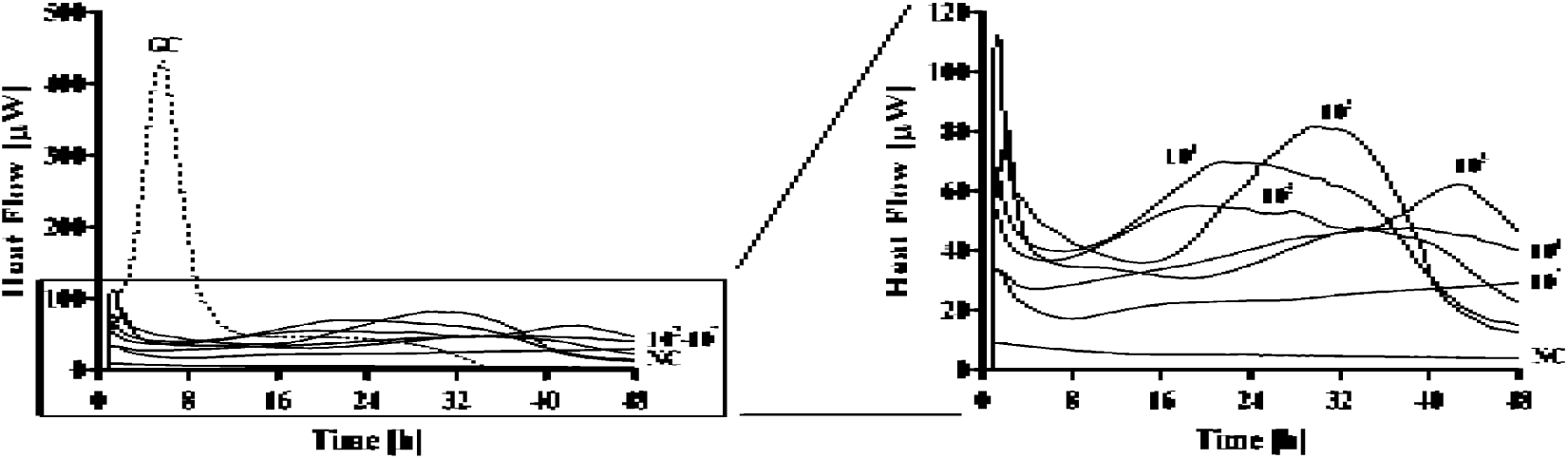
Evaluation of MRSA ATCC43300 biofilm susceptibility to Sb-1 exposure. Each curve shows the heat produced by viable bacteria present in the biofilm after 24 hour-treatment with different phage titers (ranging from 10^2^ to 10^7^ PFU/mL). Numbers above curves represent Sb-1 titers (in PFU/mL). GC, growth control (dashed line); NC, negative control.

**Table 1.**
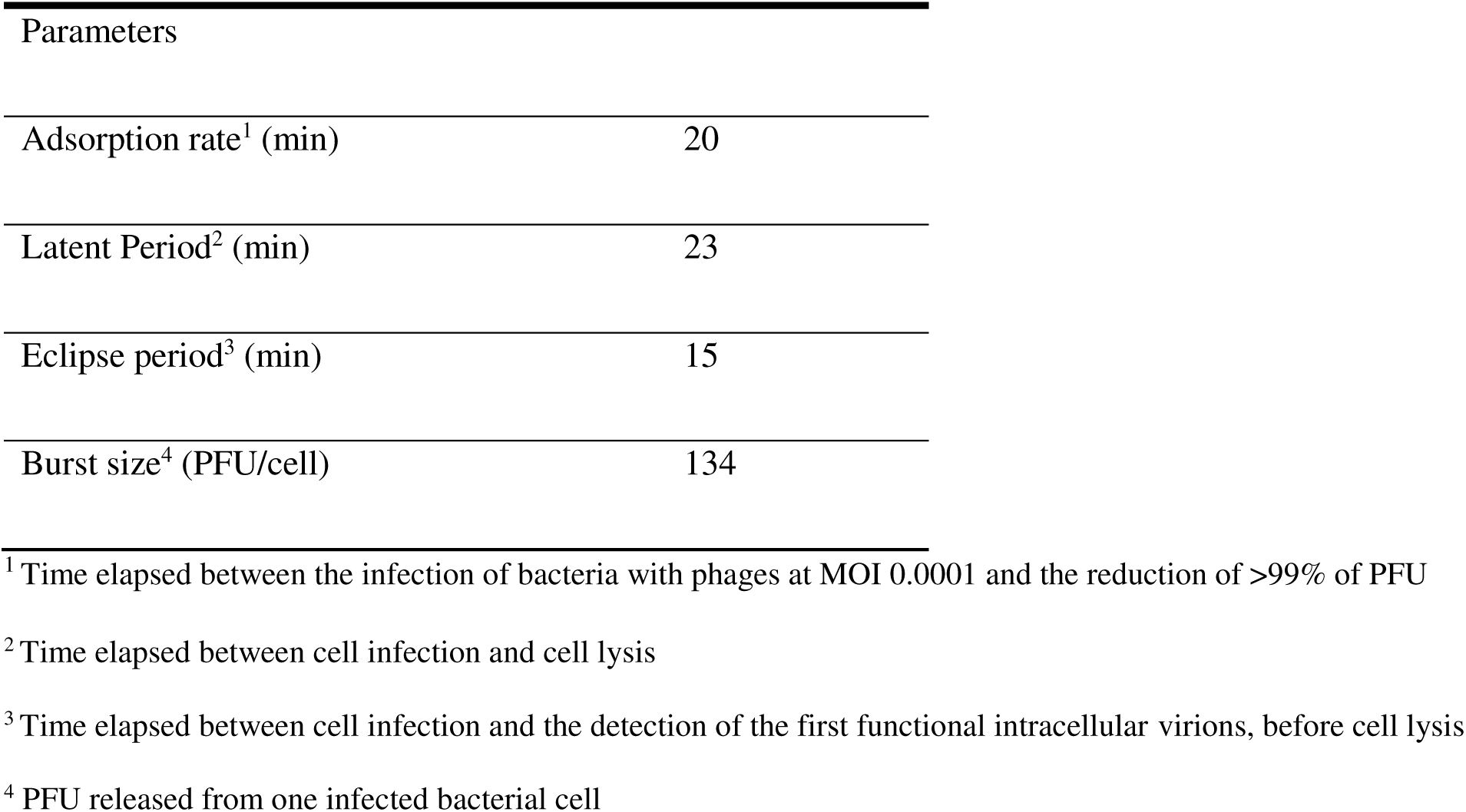
Parameters of the viral cycle of phage Sb-1.

### Simultaneous treatment with Sb-1 and rifampin/daptomycin results in eradication of *S. aureus* biofilm

We evaluated whether simultaneous treatment with Sb-1 and conventional antibiotics may produce a synergistic effect resulting in the complete eradication of the biofilm. We first tested the susceptibility of both planktonic and biofilm-embedded *S. aureus* ATCC 43300 cells to five antibiotics characterized by different mechanism of action (Figures 2 and 3). The results are shown in Table 2. As expected, although planktonic cells were susceptible to all the antibiotics tested, their sessile counterparts were resistant. However, when sub-eradicating concentrations of antibiotics together with sub-eradicating titers of Sb-1 (10^4^ and 10^5^ PFU/mL) were simultaneously used to treat biofilm-coated beads, a delay and/or reduction of heat flow produced by bacteria was observed for all the antibiotics tested (Figure 4 and Table 3), although the effect was minimal in the case of fosfomycin (Figure 4A). Notably, a synergistic effect occurred when using 10^5^ PFU/mL Sb-1 in combination with either 64 μg/ml rifampin (Figure 4B) or 32 μg/ml daptomycin (Figure 4D), which resulted in the complete eradication of the treated biofilm, as attested by the absence of growth on plate after sonication of the beads. No synergistic effects were observed in the case of fosfomycin and vancomycin (Figure 4, A and C).

**Figure 2.**
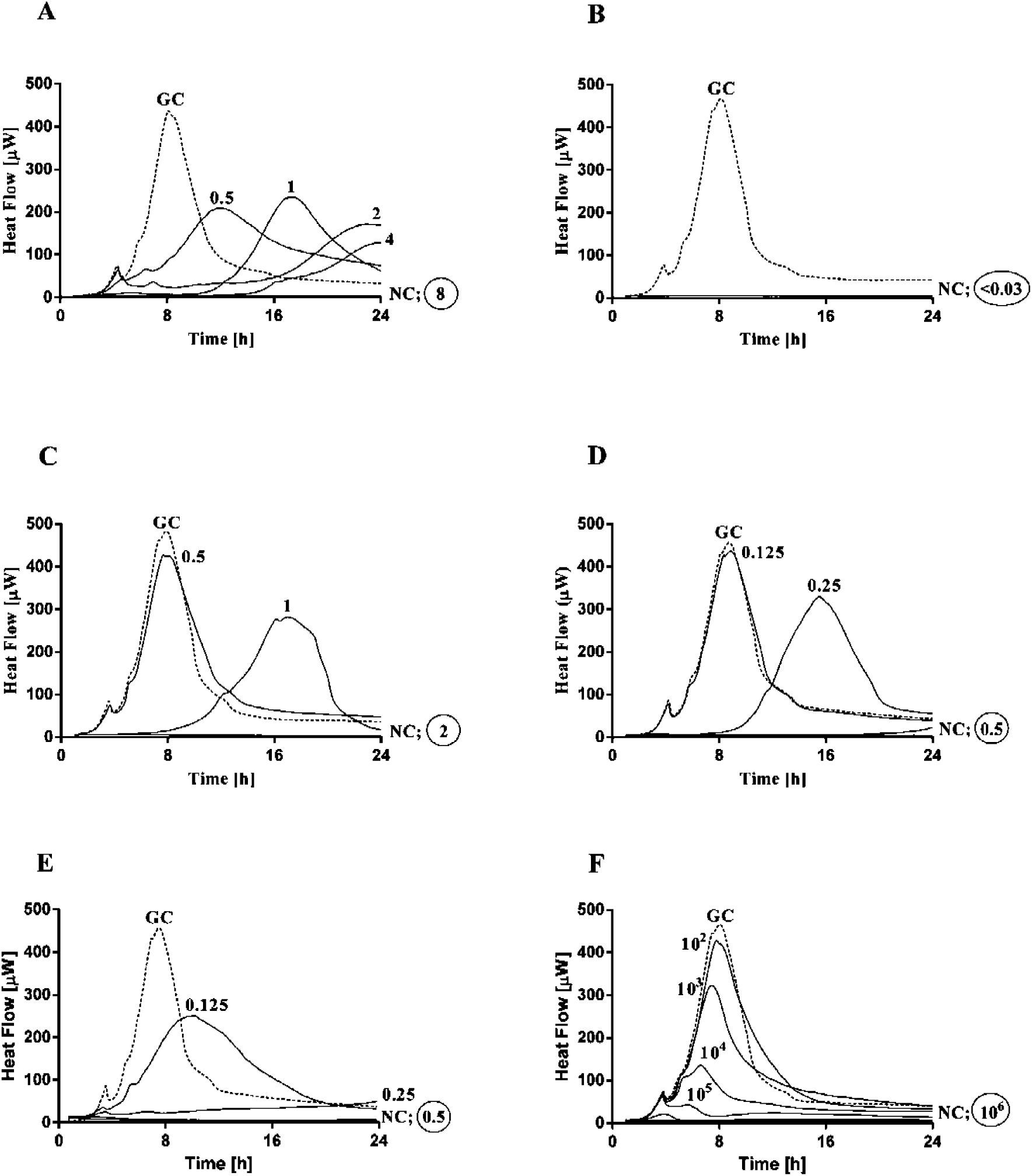
Evaluation of planktonic MRSA ATCC43300 susceptibility to fosfomycin (A), rifampicin (B), vancomycin, (C) daptomycin (D) and ciprofloxacin (E) and Sb-1 (F) exposure by isothermal microcalorimetry. Each curve shows the heat produced by viable bacteria (10^6^ CFU/ml) during the incubation with different antibiotic concentration (μg/ml) or phage titers (in PFU/mL) represented by the numbers above curves. Circled values represent the MHIC, defined as the lowest antimicrobial concentration inhibiting growth-related heat production after 24 h GC, growth control (dashed line); NC, negative control.

**Figure 3.**
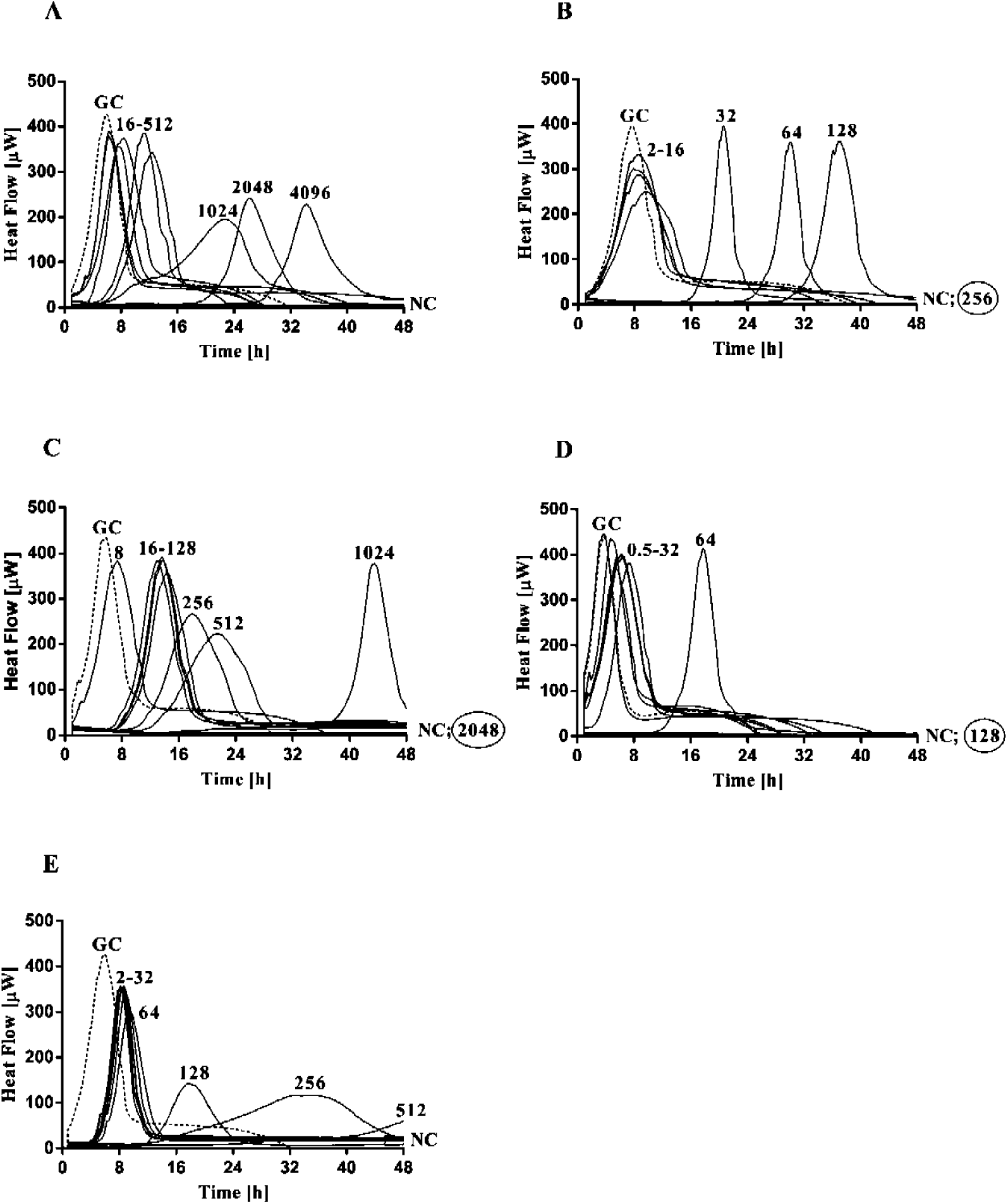
Evaluation of biofilm MRSA ATCC43300 susceptibility to fosfomycin (A), rifampicin (B), vancomycin, (C) daptomycin (D) and ciprofloxacin (E) by isothermal microcalorimetry. Each curve shows the heat produced by viable bacteria present in the biofilm after 24h of antibiotic treatment or no treatment. Numbers represent different antibiotic concentrations (in μg/ml). Circled values represent the MBEC, defined as the lowest antimicrobial concentration leading to absence of bacterial regrowth after 48 h and no colonies after sonication and plating. GC, growth control (dashed line); NC, negative control.

**Figure 4.**
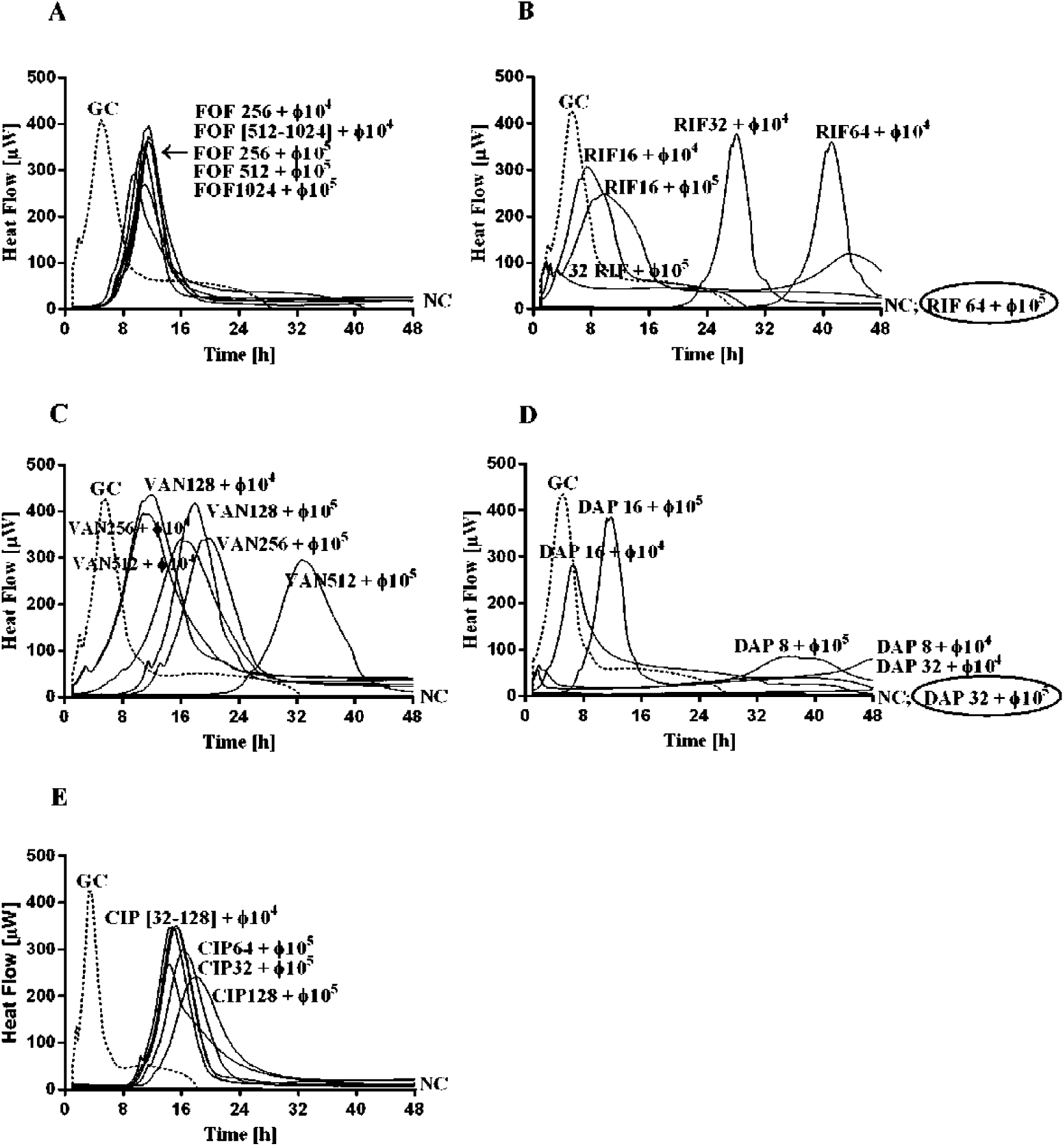
Evaluation of MRSA ATCC43300 biofilm to simultaneous exposure of Sb-1 together with fosfomycin (A), rifampin (B), vancomycin, (C) daptomycin (D) and ciprofloxacin (E) by microcalorimetry. Each curve shows the heat produced by viable bacteria present in the biofilm after 24h treatment with an antibiotic and the phage. The combinations were tested with fixed concentrations of antibiotics (1/4, 1/8 and 1/16 × the MBEC_biofilm_) and either 10^4^ or 10^5^ PFU/mL (sub-inhibitory titers) of phage Sb-1. Numbers above curves represent antibiotic concentrations (in μg/ml) and titers of Sb-1 (in PFU/mL). Circled values represent the MBEC, defined as the lowest antimicrobial concentration leading to absence of bacterial regrowth after 48 hours and no colonies after sonication and plating. GC, growth control (dashed line); NC, negative control.

**Table 2.** Antimicrobial susceptibility of planktonic and biofilm-embedded cells of *S. aureus* (ATCC 43300)

|  | Planktonic | Biofilm |  |
| --- | --- | --- | --- |
|  | MHIC<sup>2</sup> | MBBC<sup>3</sup> | MBEC<sup>4</sup> |
| **FOF<sup>1</sup> (µg/ml)** | 8 | >4096 | >4096 |
| **RIF<sup>1</sup> (µg/ml)** | <0.03 | 256 | 256 |
| **VAN<sup>1</sup> (µg/ml)** | 2 | 2048 | 2048 |
| **DAP<sup>1</sup> (µg/ml)** | 0.5 | 128 | 128 |
| **CIP<sup>1</sup> (µg/ml)** | 0.5 | >256 | >256 |
| **Sb-1 (PFU/mL)** | 10<sup>6</sup> | 10<sup>7</sup> | >10<sup>7</sup> |

<sup>1</sup> FOF: fosfomycin, RIF: rifampin, VAN: vancomycin, DAP: daptomycin, CIP: ciprofloxacin
<sup>2</sup> MHIC, minimal heat inhibitory concentration
<sup>3</sup> MBBC, minimum biofilm bactericidal concentration
<sup>4</sup> MBEC, minimum biofilm eradicating concentration

**Table 3.**
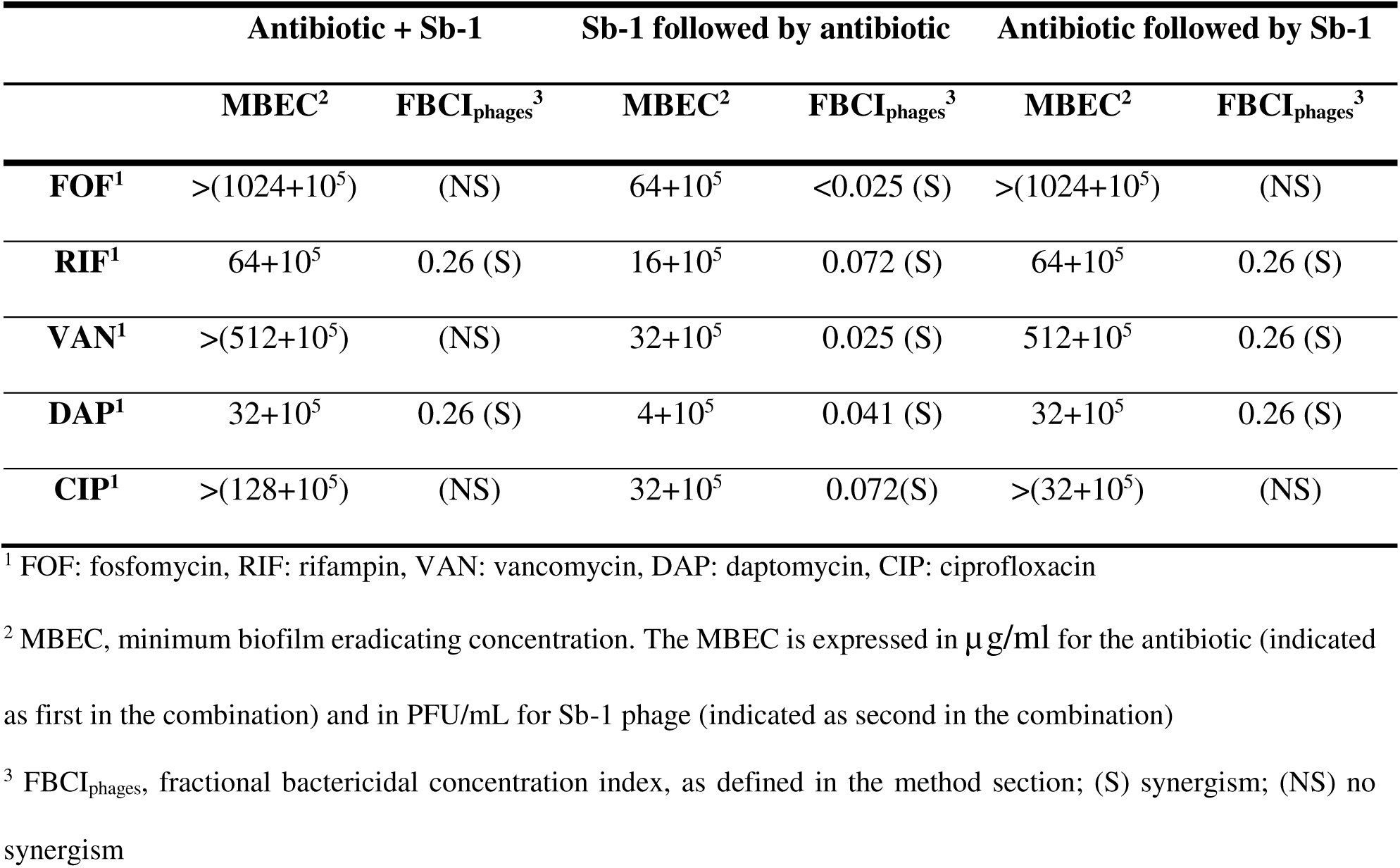
MBEC and FBCI_phage_s of antibiotic-phage combinations against *S. aureus* (ATCC 43300^)^ biofilms, as a result of either simultaneous or staggered exposure to the antimicrobial combination.

### Staggered phage and antibiotic treatment is the most effective for biofilm eradication

We then tested the effect of uncoupling the phage/antibiotic treatment and replacing it with a staggered treatment. We treated the biofilm-coated beads for 24 hours with sub-eradicating amount of agent A (either Sb-1 or antibiotic), removed it by washing, and incubated for further 24 hours with sub-eradicating amount of agent B (either antibiotic or Sb-1). The viability of the cells still adhering to the beads was assessed by microcalorimetry. Interestingly, the staggered phage – antibiotic treatment more pronouncedly inhibited the heat flow production for all the combinations tested, as compared with the simultaneous treatment. In particular, the pre-treatment with 105 PFU/mL Sb-1 resulted in a synergistic eradicating effect with all the antibiotics tested, including fosfomycin and vancomycin (Figure 5 and Table 3). Moreover, the synergistic eradicating concentrations of rifampin and daptomycin were 2 and 3 dilutions below the ones in the simultaneous treatment, respectively. A complete inhibition of the heat flow production, although not eradicating, was also induced by the pre-treatment with 10^4^ PFU/mL Sb-1 followed by incubation with even lower antibiotic concentrations (Figure 5, fosfomycin ≤ 16 μg/ml, rifampin ≤ μg/ml, vancomycin 32 μg/ml, and daptomycin ≤ 0.5 μg/ml). A different scenario was observed for the staggered antibiotic – phage treatment. Firstly, no synergistic effect could be observed in the case of fosfomycin (Figure 5). A synergistic effect in the case of the pre-treatment with rifampin, vancomycin, and daptomycin followed by incubation with 10^5^ PFU/mL Sb-1 could only be observed at higher antibiotic concentrations (Figure5).

**Figure 5.**
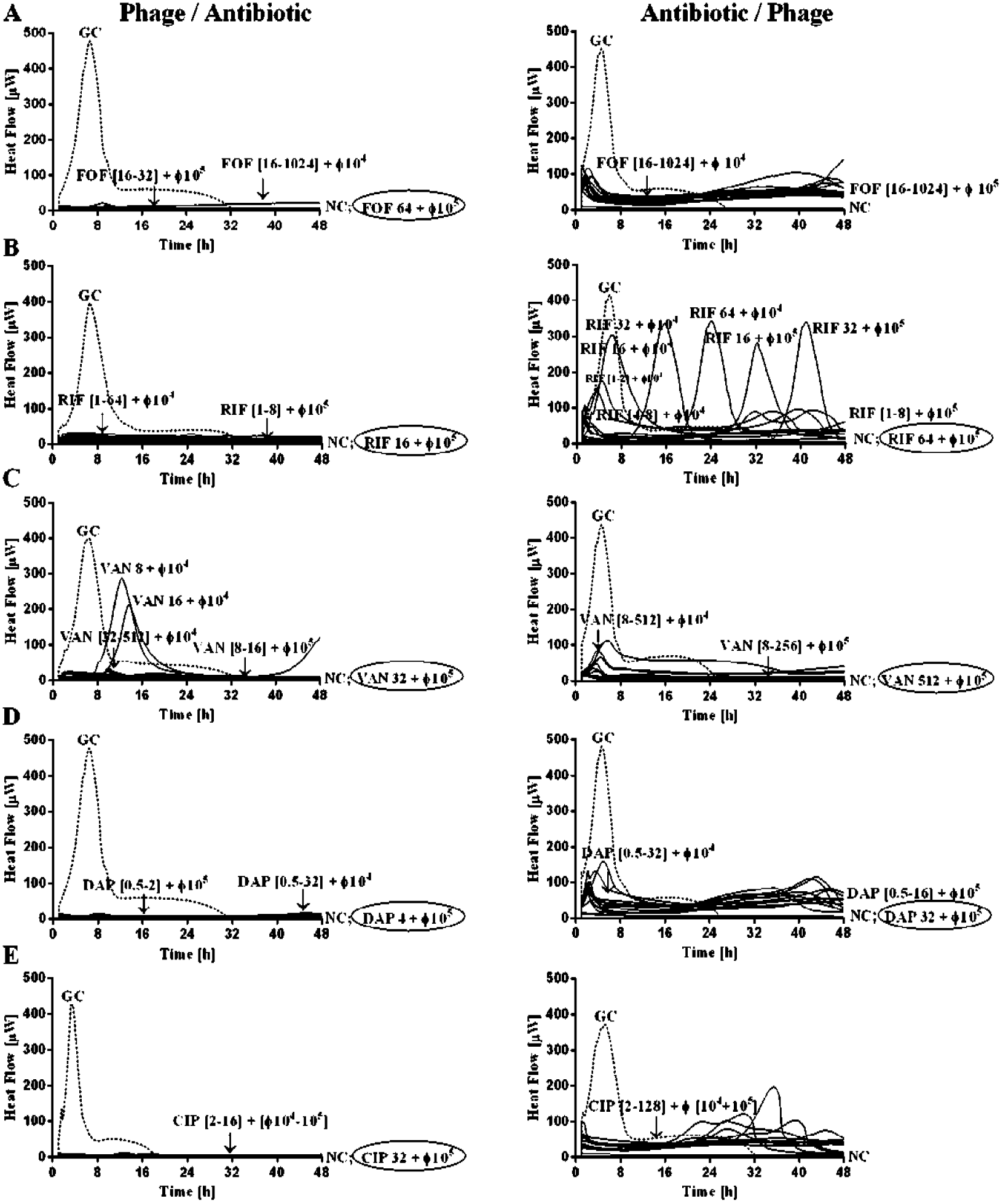
Evaluation of MRSA ATCC43300 biofilm to staggered exposure of Sb-1 followed (graphs on the left) or preceded (graphs on the right) by fosfomycin (A), rifampin (B), vancomycin, (C) daptomycin (D) and ciprofloxacin (E) by microcalorimetry. Each curve shows the heat produced by viable bacteria present in the biofilm after the staggered treatment. Untreated controls were also added. The combinations were tested with fixed concentrations of antibiotics (1/4, 1/8, 1/16, 1/32, 1/64, 1/128 and 1/256 × the MBEC biofilm) and either 10^4^ or 10^5^ PFU/mL (sub-inhibitory titers) of phage Sb-1. Numbers above curves represent antibiotic concentrations (in μg/ml) and titers of Sb-1 (in PFU/mL). Circled values represent the MBEC, defined as the lowest antimicrobial concentration leading to absence of bacterial regrowth after 48 hours and no colonies after sonication and plating. GC, growth control (dashed line); NC, negative control.

### Sb-1 degrades the extracellular polysaccharide matrix

The effect of Sb-1 on biofilm matrix of MRSA ATCC43300 was assessed by confocal microscopy. A 24 hours old biofilm was stained with two dyes, specific for the poly-N-acetylglucosamine residues of the extracellular polysaccharides and the cellular DNA, respectively. As we expected, the polysaccharide component was clearly visible in the untreated control. Treatment of the biofilm with increasing – but sub-eradicating – titers of Sb-1 resulted in the progressive, degradation of the polysaccharide component (Figure 6), as attested by the statistically significant reduction of fluorescence mean intensity (Figure 7), although it did not affect cell viability (Figure 8).

**Figure 6.**
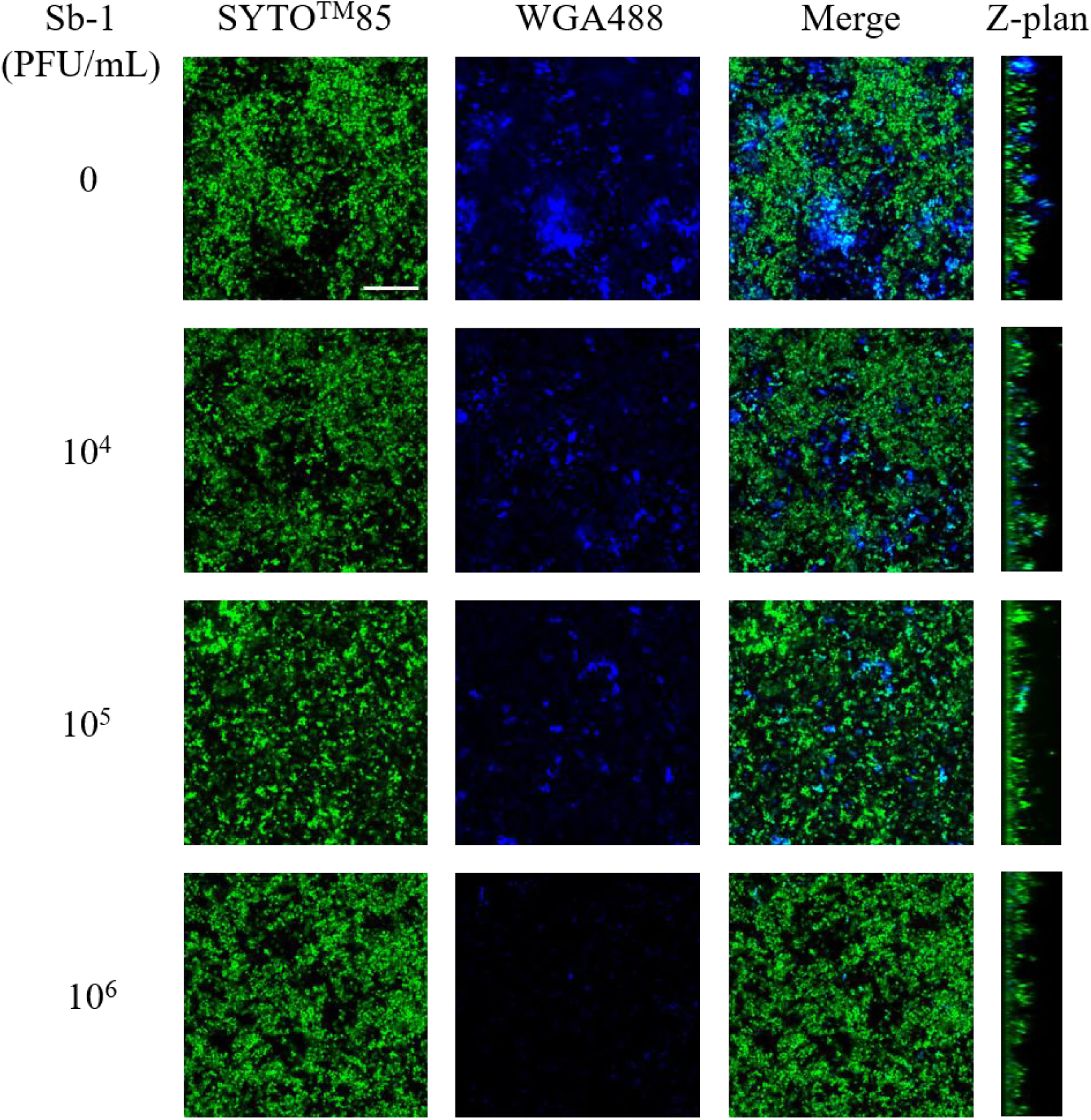
CLSM images of MRSA ATCC43300 biofilm untreated and treated with Sb-1. MRSA biofilm (24h-old) was exposed for 24 hours to different Sb-1 titers (ranging from 10^4^ to 10^6^ PFU/mL) and then stained with green fluorescent labeled WGA488 (488/500–600 nm) for exopolysaccharides and SYTO^™^85 (561/600–700 nm) for bacterial cells. An untreated control was also added. Scale bar: 25 μm.

**Figure 7.**
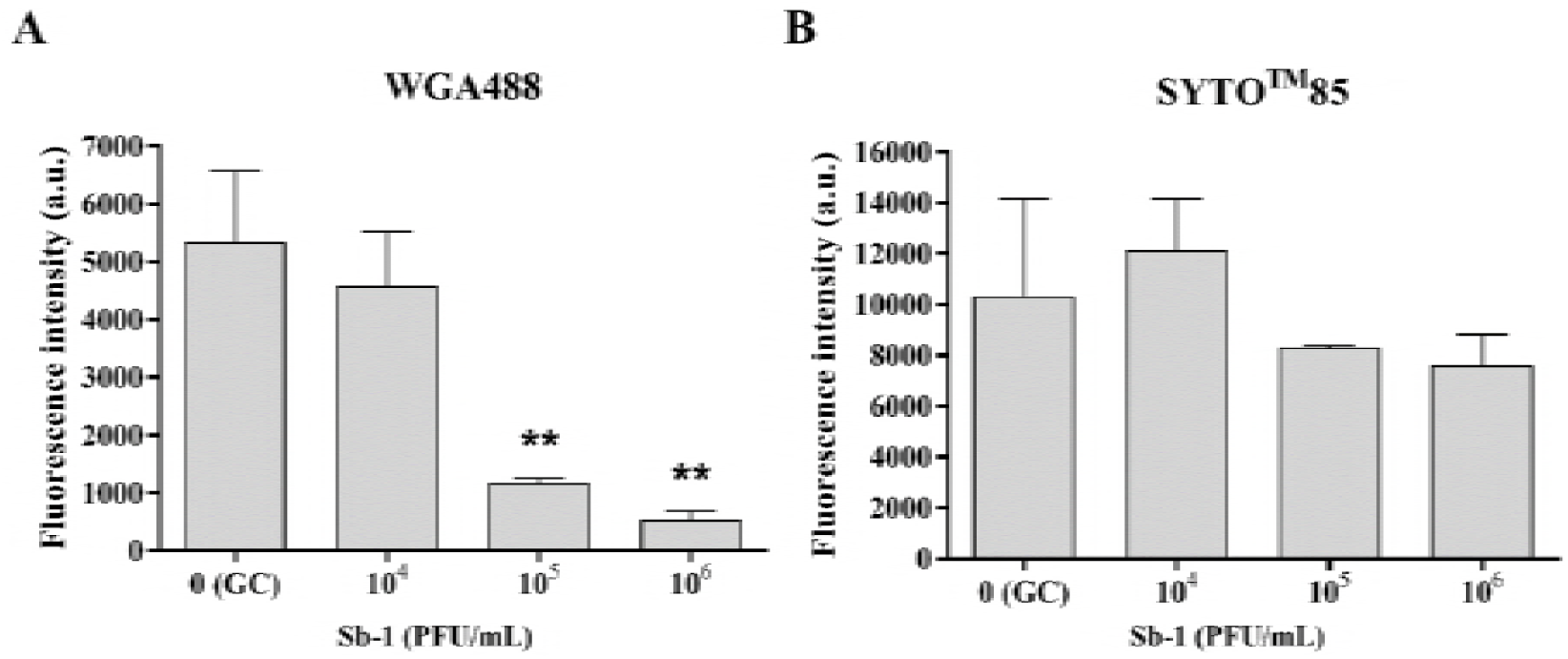
Mean fluorescence intensity of images after staining of matrix exopolysaccharides and bacteria collected by CLSM. The mean fluorescence intensity was calculated from images of biofilms treated with different Sb-1 titers (ranging from 10^4^ to 10^6^ PFU/mL) and then stained with WGA488 (A) and SYTO^™^85 (B), respectively. A region drawn outside the biofilm was used to calculate (and then subtract) the average background signal. Corrected fluorescence mean values ± SEM were expressed in arbitrary units. ^∗∗^p < 0.01; (one-way ANOVA followed by Tukey-Kramer post hoc test).

**Figure 8.**
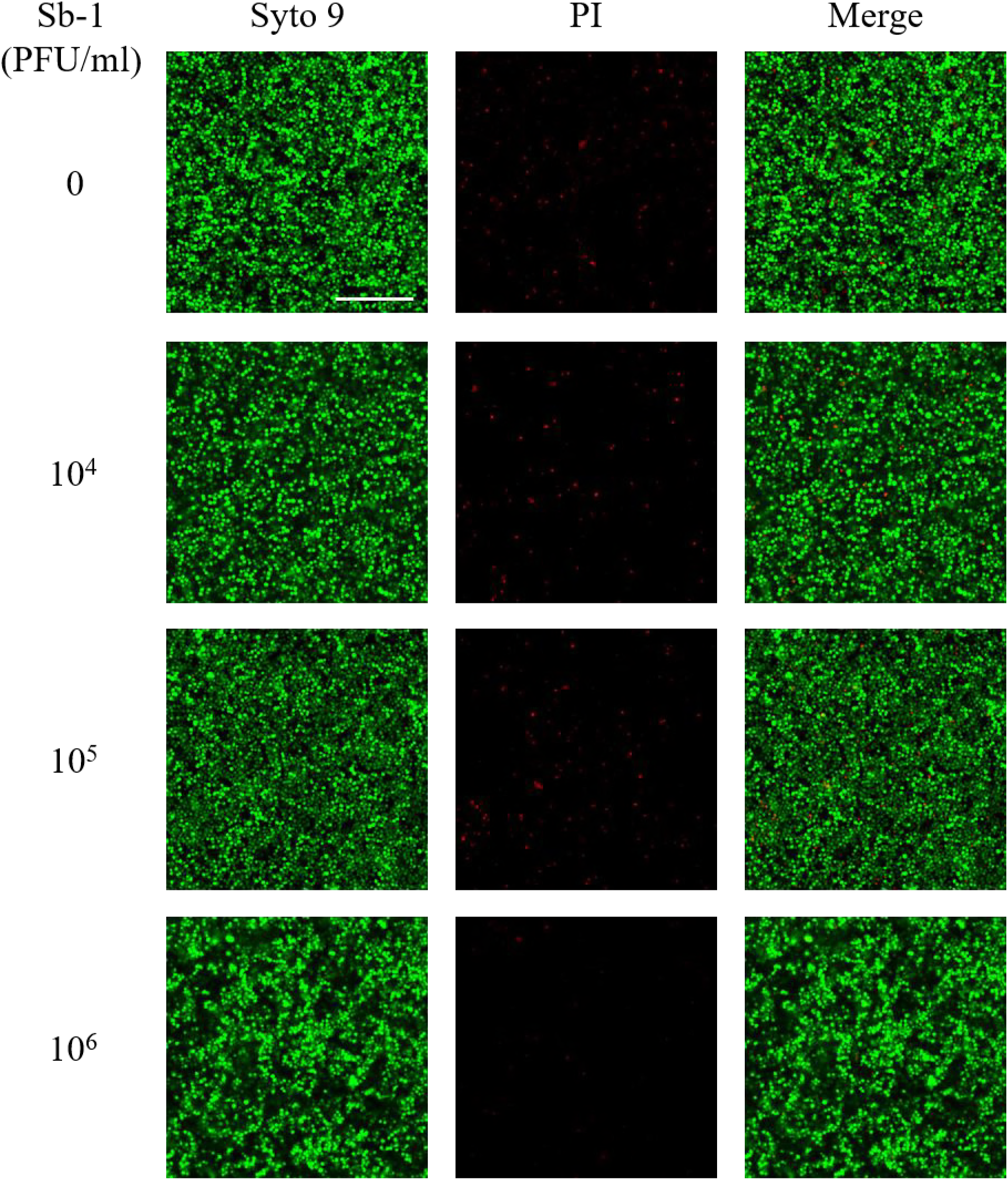
CLSM images of MRSA ATCC43300 biofilm untreated and treated with Sb-1. MRSA biofilm (24h-old) was exposed for 24 hours to different Sb-1 titers (ranging from 10^4^ to 10^6^ PFU/mL). The viability of the cells was evaluated stained with green fluorescent labeled Syto 9 (488/500–540 nm) for alive bacteria and with red fluorescent Propidium iodide (PI) (561/600–650 nm) for dead bacteria. An untreated control (0) was also added. Scale bar: 25 μm.

### Sb-1 targets also persister cells

We tested the activity of Sb-1 on free floating i) cells scraped from the biofilm; ii) ciprofloxacin-selected persister cells scraped from the biofilm; iii) stationary phase cells; iv) CCCP-induced persister-like cells. The instantaneous cell viability after phage treatment was evaluated by CFU counting. As shown in Figure 9 no viability reduction was observed following treatment with 10^4^ PFU/mL in any of the conditions tested. By contrast, the 10^7^ PFU/mL titer determined a reduction of the CFU/mL (≈ 2-5 log 10 CFU), demonstrating that the phage displayed lytic activity regardless of the metabolic state of the cells. Interestingly, despite the fact that neither titer completely eradicated the bacteria, when the treated cells from each condition were inoculated into fresh media, no growth was observed over the course of 24 hours (Figure 9). No colonies were observed when these cultures were plated. Indeed, the reversion of *S. aureus* cells pre-treated with sub-eradicating titers of Sb-1 to a metabolically active state resulted in the complete killing of the bacterial population.

**Figure 9.**
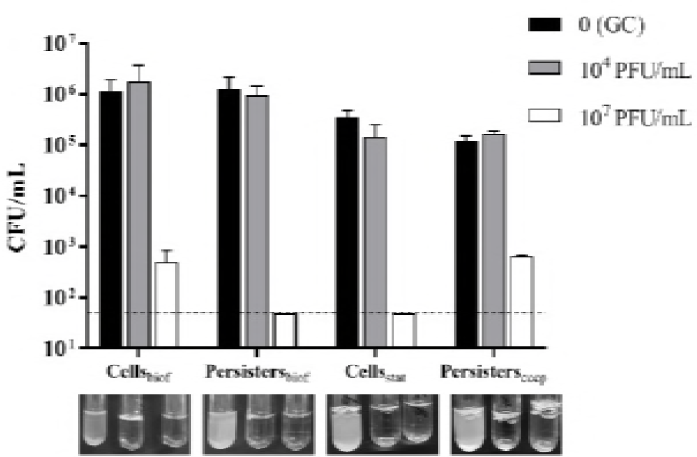
Lytic effect of Sb-1 against free-floating bacterial cells of MRSA ATCC43300 in different metabolic conditions. Cells detached from biofilm (Cells_biof_), persister cells isolated from biofilm after treatment with ciprofloxacin (Persisters_biof_), stationary phase cells (Cells_stat_) and CCCP-induced persisters (Persisters_cccp_) were exposed to either 10^4^ and 10^7^ PFU/mL Sb-1 in PBS+1% BHI for 3 hours at 37°C. Untreated controls were also added. After phage treatment, cells were plated to enumerate CFUs. Data are means with standard deviation, and the dotted line represents the detection threshold. A set of samples treated as above-mentioned and washed to remove un-adsorbed phages, was also inoculated in fresh BHI for 24 hours and then the turbidity was visually evaluated (on the bottom panel).

## DISCUSSION

Biofilm-associated infections are difficult to eradicate due to their tolerance and refractivity to conventional antibiotics (17). As a result, bacteriophages have been regaining interest as potential alternative strategy for biofilm treatment (18-20). Sb-1 is a *S. aureus* specific virulent phage active versus both antibiotic-susceptible and antibiotic-resistant strains, with a low percentage of bacterial strains (<10%) showing resistance to its infection (21). This trend was confirmed, as Sb-1 exhibited a broad lytic spectrum against both MRSA and MSSA strains isolated in implant-associated infections (>90% MRSA strains susceptible). Microcalorimetric analysis indicated that 10^7^ PFU/mL Sb-1 could exert killing activity versus ATCC 43300 biofilm, but it did not result in a complete eradication of the biofilm, as attested by plating. These results are in agreement with *in vivo* data obtained in a rat implant-related infection model (22). It is now generally accepted that a single-dose phage application to even single-strain laboratory biofilms can result in incomplete elimination of bacteria (23, 24), possibly due to the presence of the matrix and non-metabolically active cells (25). In this scenario, the combination of antibiotic and bacteriophage therapy might represent an alternative strategy for pursuing the clearance of biofilm-embedded cells. Simultaneous treatment with Sb-1 and either rifampin or daptomycin resulted in a synergic eradication of ATCC 43300 biofilm. To the best of our knowledge, for the first time the synergism between phages and daptomycin against biofilm was evaluated. Few studies investigated the effect of phage-derivative enzymes in combination with daptomycin, but they focused on the survival from bacteremia in a mice model, in which biofilm embedded cells are most likely not involved (26, 27). A strong synergistic effect of a phage-rifampin combination on MRSA biofilm has also been observed by Rahman et al (28). Although rifampin remains the only antibiotic that shows high efficacy against biofilm-associated staphylococci (29, 30), bacteria are prone to easily develop resistance to this compound (31, 32), and therefore its use in combination with phages may result in the viral particles targeting rifampin-resistant bacteria. We also investigated the effect of staggered treatment, and we observed that the exposure to the phage followed by antibiotic treatment is the most effective delivery scheme for biofilm eradication. In the case of rifampin and daptomycin, in which the co-incubation with Sb-1 already determined a synergistic effect, the pre-exposure to Sb-1 followed by antibiotic treatment completely eradicated the MRSA biofilm by even lower antibiotic concentrations. Interestingly, the treatment of the cells with Sb-1 before being exposed to fosfomycin, vancomycin, and ciprofloxacin resulted in a synergistic effect that could not be observed in the simultaneous treatment. In fact, despite the fact that the simultaneous administration of Sb-1 and fosfomycin (up to 1024 μg/ml) had no effect on MRSA biofilm, a 10^5^ PFU/mL phage pre-treatment strongly enhanced the bactericidal activity of the antibiotic. In this condition a fosfomycin concentration still reachable in the clinical practice completely eradicated MRSA biofilm. A similar effect was also observed for vancomycin and ciprofloxacin. The promising potential of the phage-antibiotic staggered treatment has been recently highlighted by other two groups (14, 15), and may be related to the potential antagonistic effect of simultaneous treatment with antibiotics, which inhibit and/or kill bacteria, and bacteriophages, which require replicating bacterial cells to propagate (33, 34).

A synergistic effect resulting in a strong killing was observed for low concentrations of all the antibiotics tested when preceded by the lowest phage titer tested (10^4^ PFU/mL). Although in our *in vitro* model these combinations were not eradicating, this strong suppressive effect may have a relevance in the context of a human infection, where a complete eradication may be obtained with the contribution of the host immune system. Our microscopy observations showed that sub-inhibiting titers of Sb-1 were able to degrade the extracellular polysaccharide component of the matrix of ATCC 43300, most likely thanks to the presence of degrading tail enzymes. As it is known that the diffusion of some antibiotics may be blocked/delayed by *S. aureus* matrix (35, 36), it is plausible that the degradation of some of its components may facilitate their diffusion to the bottom of the sessile community.

We also tested Sb-1 activity against both less-metabolically active cells and persister cells, which greatly contribute to the tolerance of biofilm to conventional antibiotics. 10^4^ PFU/mL Sb-1 did not reduce at all bacterial viability, which was not surprising, as phages need a metabolically active host for their replication.

However, we found that 10^7^ PFU/mL Sb-1 strongly reduced bacterial viability. This may be explained with a “lysis from without” (37): high phage/bacteria ratio may result in a direct bacterial lysis that is not dependent on viral replication and phage production, and so not affected by the metabolic state of bacteria. Notably, when persister cells infected with the lowest phage titer reverted to a normal-growing phenotype no CFU could be observed after 24 hours. This may be due the activation of phage progeny formation from the previously internalized viral nucleic acid. As soon as the bacterial cell reactivates its metabolism, this “*trojan horse effect*” results in bacterial cell lysis. This may have a clinical impact, as using phages in combination with conventional antibiotics may help avoiding the relapsing of the infection due to the recalcitrance of persister cells.

In conclusion, this work provides insights into Sb-1 data and possible explanations regarding the phage/antibiotic exposure to eradicate *S. aureus* biofilm. An *in vivo* assessment may generate further insights and support the development of phage/antibiotic combination therapy.

## MATERIALS AND METHODS

### Bacteria and phages

*S. aureus* ATCC 43300 and 57 *S. aureus* clinical strains isolated from patients with orthopaedic implant-associated infections were used for this study. Purified Sb-1 bacteriophage was supplied by Georgia Eliava Institute (Tbilisi, Georgia). The susceptibility of 28 methicillin-resistant *S. aureus* (MRSA) and 29 methicillin-sensitive *S. aureus* (MSSA) strains to Sb-1 was evaluated by spot test (38).

### Evaluation of Sb-1 infectious parameters

The infectious parameters were determined using the ATCC 43300 S. aureus strain. Sb-1 adsorption rate was determined as in (28). Sb-1 growth cycle parameters were evaluated by performing a one-step growth experiment as in (39)

### Antimicrobial assay

The analyses were performed using an isothermal calorimeter (TAM III; TA Instruments, USA). The evaluation of the antimicrobial activity of either the antibiotics or Sb-1 versus either planktonic or biofilm-embedded cells of ATCC 43300 *S. aureus* was performed as previously described (16, 40), with minor modifications. These modifications included that i) the bacterial cells were grown in BHI broth; ii) for the anti-biofilm tests, 2-3 colonies of ATCC 43300 were inoculated into BHI broth and incubated in the presence of glass beads at 37°C for 24 hours. The minimum heat inhibitory concentration (MHIC) for planktonic bacteria was defined as the lowest antimicrobial concentration inhibiting growth-related heat production after 24 hours. The minimum biofilm bactericidal concentration (MBBC) measured by calorimetry was defined as the lowest antimicrobial concentration leading to lack of heat production related to the absence of bacterial re-growth after 48 hours of incubation in the microcalorimeter (16). The minimum biofilm eradicating concentration (MBEC) of antibiotic combinations, defined as the lowest concentration of antibiotic required to eradicate the biofilm (0 CFU/bead on plate counts) (16), was evaluated by CFU counting of the sonicated beads.

In order to evaluate the effect of combined treatment (antibiotic + Sb-1) on biofilm-embedded cells, mature biofilms were grown on the beads as described above, washed, and then incubated in BHI broth in the presence of sub-eradicating concentrations/titers of a given antibiotic and Sb-1, respectively, for 24 hours at 37°C. The effect of sequential 24 hour-exposure to Sb-1 followed by a 24 hour-exposure to a given antibiotic (and *vice versa*) was also evaluated. In all cases, the heat flow produced by the viable cells still embedded in the biofilm was measured for 48 hours by calorimetric analysis.

We adapted the commonly used fractional inhibitory concentration index (FICI) formula to take into account that i) Sb-1 titers are evaluated in 10-fold serial dilutions ii) we were interested in combinations capable of eradicating the biofilm. The fractional bactericidal concentration index (FBCI_phages_) was defined based on the MBECs of the individually active antimicrobial agents A and B as: (MBEC _A in combination_/MBEC_A alone_) + MBEC_B in combination_/MBEC_B alone_. If the FBCI_phages_ is ≤ 0.26, the concentrations of agent A and B used in the combination have a synergistic effect that results in the complete eradication of the biofilm. If the FBCI_phages_ is > 0.26, there is no synergistic effect.

### Confocal Laser Scanning Microscopy (CLSM)

The effect of Sb-1 treatment on ATCC 43300 biofilm and matrix was evaluated by CLSM. An overnight culture was diluted 1:100 and dispensed into an 8-well μ-Slide (Ibidi) in the presence of Sb-1 (10^4^ to 10^6^ PFU/mL) and incubated at 37°C for 24h. Biofilms were stained with SYTO^™^85 and Wheat Germ Agglutinin, Oregon Green^®^ 488 Conjugate (WGA488) (Life Technologies). Biofilm cell viability after Sb-1 treatment was determined by staining cells with Syto9 and propidium iodide (PI) using the LIVE/DEAD BacLight Bacterial Viability Kit (Molecular Probes, Life technologies) as recommended by the manufacturer. After staining, biofilms were washed and examined under the microscope (TCS SP5, Leica, Heidelberg, Germany) using a 63 × objective and a pinhole aperture of 1.0 Airy. WGA488 and SYTO^™^85 were excited at 488 nm and 561 nm, respectively. The following collection ranges were adopted: 500–540 nm (WGA488) and 600–700 nm (SYTO™85). For viability assay, samples were sequentially excited at 488 (Syto9) and 561 (PI) nm and emissions were monitored at 500–540 and 600–650 nm, respectively. For each image, the mean of fluorescent intensity was calculated as previously described (41).

### Evaluation of Sb-1 activity against CCCP-induced and ciprofloxacin-induced persister cells

Persister status was induced in ATCC 43300 bacteria following two different methods. Firstly, persister cells were induced using carbonyl cyanide m-chlorophenylhydrazone (CCCP, Sigma-Aldrich) as reported in (42). Alternatively, a 24 hours-old biofilm was treated with 512 μg/ml ciprofloxacin (or PBS as control) for 24 hours at 37°C. At the end of the treatment with either CCCP or ciprofloxacin, cells were washed (after scraping when biofilm-embedded) and treated with 0, 10^4^, or 10^5^ PFU/mL (final concentration ≈ 5 × 10^5^ CFU/mL) for 3h at 37°C, in duplicate. For both conditions, one set of samples was then plated on BHI agar for CFU count, and the other was inoculated in fresh BHI broth and incubated for 24 hours at 37°C before being visually assessed for growth and plated for CFU counting.

### Statistical analysis

Differences between the mean values of groups were evaluated by one-way analysis of variance (ANOVA) followed by Tukey-Kramer test. Data presented represent the mean ± the SEM of at least three independent experiments. A p value <0.05 was considered statistically significant. The graphs in the figures were plotted using Prism software (version 6.01; GraphPad Software, La Jolla, CA).

## ACKNOWLEDGMENTS

This work was supported by PRO-IMPLANT Foundation (www.proimplant-foundation.org) in Berlin.

